# Calibration of MRI-based reference intervals to new samples

**DOI:** 10.1101/2025.11.25.690453

**Authors:** Andrew A. Chen, Jakob Seidlitz, Margaret Gardner, Richard A. I. Bethlehem, Lena Dorfschmidt, Eren Kafadar, Lifespan Brain Chart Consortium, Andreana Benitez, Jens H. Jensen, Simon Vandekar, Theodore D. Satterthwaite, Aaron F. Alexander-Bloch

## Abstract

Reference intervals, defined as intervals containing a new observation with a specified probability relative to reference data, would be clinically useful in assessing brain magnetic resonance imaging (MRI). Brain charts, which are estimates of MRI phenotypes across covariates such as age and sex, can be used to construct reference intervals. However, the reference data used to fit intervals often differs from a new sample in terms of study design, MRI acquisition, and image preprocessing. Application of MRI reference intervals to new samples remains a challenging problem. Here, we propose a new method called Reference interval calibration via conFormal prediction (ReForm) that adjusts reference intervals for a new sample. Our method builds on recent work in conformal prediction, which yields intervals with guaranteed coverage for new observations. Through resampling experiments in Lifespan Brain Chart Consortium cortical thickness data, we compare ReFormed reference intervals to refitting intervals, statistical harmonization methods, and model-based adjustment of intervals. Notably for patient privacy concerns, ReForm does not require sharing of reference data. Yet, our empirical results demonstrate that ReForm controls FPR similarly or better than alternative methods which require sharing reference data. Finally, we provide recommendations for practical applications of ReForm and an R package (https://github.com/andy1764/ReForm) for calibrating reference intervals using ReForm.

## 1 Introduction

Imaging consortia increasingly aggregate massive databases of magnetic resonance imaging (MRI) scans for precise mapping of the brain across the lifespan. Their goal is to estimate *brain charts*for a given MRI phenotype, which are models that capture nonlinear trajectories across neurodevelopment and aging while accounting for known sex differences and other covariates (Bethlehem et al., 2022; Dima et al., 2022; Frangou et al., 2022; Rutherford et al., 2022). Numerous consortia have recently published brain charts across additional types of MRI scans or open-access resources for other researchers to fit their own brain charts (Kim et al., 2025; Shafiei et al., 2025; Zhu et al., 2025). The promise is that brain charts can enable scientists and clinicians to compare new scans to a well-characterized healthy reference population.

However, the promise of brain charts is limited due to differences in MRI acquisition and preprocessing. Differences across MRI acquisition have been recognized to introduce bias and reduce power (Han et al., 2006) and several recent studies highlight this issue across multiple MRI modalities (Cetin Karayumak et al., 2019; Fortin et al., 2017, 2018; Shinohara et al., 2017; Yu et al., 2018). Simultaneously, several groups have acknowledged that different preprocessing steps can lead to substantially different results across a wide range of MRI modalities and derived phenotypes (Antonopoulos et al., 2023; Bhagwat et al., 2021; Botvinik-Nezer et al., 2020; Haddad et al., 2019).

Recent work in brain charts has led to several statistical approaches to dealing with technical biases resulting from heterogeneity in MRI acquisition and preprocessing. One category is *harmonization* methods, which aim to transform data and eliminate systematic biases attributable to acquisition and preprocessing differences (Fortin et al., 2017, 2018; Gardner et al., 2025; Hu et al., 2023; Johnson et al., 2007; Ling et al., 2022). Another category is *model adjustment* which aims to fit model-specific adjustment parameters in new data to account for differences across study sites. Model adjustments are tailored to a specific choice of model used to fit the brain chart, typically a generalized additive model for location, scale, and shape (GAMLSS; Rigby and Stasinopoulos, 2005). Both categories of methods have found widespread use across a number of recent brain chart studies (Bethlehem et al., 2022; Gardner et al., 2025; Ge et al., 2024; Kim et al., 2025; Shafiei et al., 2025; Zhu et al., 2025).

In clinical chemistry, healthy populations are typically used to construct reference intervals that describe the range of normal values for a given phenotype (Gräsbeck and Fellman, 1968; Ozarda, 2016). The definition of reference intervals requires that an observation from the reference data falls inside the interval with a specified probability. Practically, the Clinical and Laboratory Standards Institute (CLSI) EP28-A3c guidelines adopted in 2008 recommend verifying reference intervals by taking a new sample with a specified tolerance for the proportion of samples falling outside the interval (Clinical and Laboratory Standards Institute, 2008), which we refer to as the false positive rate (FPR). Inspired by these guidelines, we propose a stricter definition for a valid reference interval, requiring a new observation to fall above and below the interval with a specified probability.

Brain charts, which are an estimate of the distribution of MRI phenotypes conditional on age and sex, can be used to construct reference intervals. Several studies publish their brain charts or reference intervals for other researchers to utilize (Bethlehem et al., 2022; Kim et al., 2025; Zhu et al., 2025). However, MRI reference intervals currently offer no guarantees of FPR control for a new sample. Neither harmonization nor model adjustment methods have been tested for constructing reference intervals that control FPR.

Here, we draw from recent work in conformal prediction (Lei et al., 2018; Romano et al., 2019b) to propose a method for adjusting reference intervals called Reference interval calibration via conFormal prediction (ReForm). ReForm differs from harmonization and model adjustment methods in that it guarantees control of the FPR in a new observation and thus produces valid reference intervals. Through resampling studies in the Lifespan Brain Chart Consortium (LBCC) cortical thickness data, we demonstrate the performance of ReForm compared to several alternative methods. We find that ReForm controls FPR across a number of different experiments while requiring reasonable sample sizes to achieve adequate FPR empirically. Compared to harmonization methods, we find that ReForm substantially improves FPR control while eliminating dependency on harmonization model assumptions and reference set sample size. Versus model adjustment, we find that ReForm performs similarly even without access to reference data, so long as an adequately sized calibration set is available. In Alzheimer’s disease (AD) patients, we demonstrate that our ReFormed reference intervals are sensitive to deviations in entorhinal thickness values. Finally, we offer practical recommendations for using ReForm as a way to calibrate reference intervals from any MRI phenotype for new samples.

## 2 Methods

### 2.1 Problem setting

Denote our full data (*X_i_, Y_i_*) where *X_i_* are covariates, *Y_i_* are outcomes, and *i* = 1, 2*, . . . , n*. Our full data consists of a reference set *N_R_* and *calibration set N_C_* with no overlap between the sets, such that *N_R_* ∪ *N_C_* = {1, 2*, . . . , n*} and *N_R_* ∩ *N_C_* = ∅. In neuroimaging, the reference and calibration sets often come from separate datasets with differences in study protocols, scanners, and acquisition parameters. Thus, the distribution of (*X_i_, Y_i_*) may differ between the reference and calibration sets.

Our goal is to construct *reference intervals* denoted *C*(*X*) = [*L*(*X*)*, U* (*X*)] that are *valid* at a chosen level *α*. That is, for a new sample (*X_n_*_+1_*, Y_n_*_+1_) drawn from the same dataset as the calibration set, a valid 100 × (1 − *α*)% reference interval *C*(*X*) has the property that a new observation falls above the reference interval with probability *α/*2 and below with probability *α/*2, or equivalently

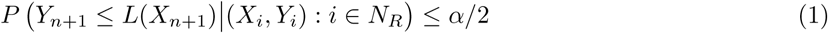

and

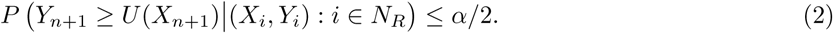

Note that this definition of validity is stricter than many standard definitions, which only consider observations within the reference dataset. Additionally, our definition is more general than common definitions of reference intervals, which often refer only to the 95% reference interval (Ozarda, 2016). Validity may not be feasible in the general setting; however, we will show that validity is achievable when the probability is taken over the calibration set and the new observation.

### 2.2 Reference interval calibration via conformal prediction (ReForm)

Here, we propose a new method that can adjust a fitted reference interval using the calibration set and produce a valid reference interval. We will begin by reviewing conformalized quantile regression, which is a type of conformal prediction method (Lei et al., 2018; Romano et al., 2019b). Suppose our full data are exchangeable and drawn from the same joint distribution such that (*X_i_, Y_i_*) ∼ *P* , *i* = 1, 2*, . . . , n*. This assumption is reasonable when the reference and calibration sets are taken from the same dataset. In this setting, conformalized quantile regression (CQR) yields a valid 100 × (1 − *α*)% reference interval. CQR first fits a quantile regression model in the reference set to construct a *naïve* reference interval *C*(*X*) = [*q̂_α/_*_2_(*X*)*, q̂*_1_*_−α/_*_2_(*X*)] where *q̂_τ_* (*X*) is the estimated conditional *τ* -th quantile function of *Y* given *X*. Here, we focus on the version of CQR which controls the upper and lower coverage of the interval. Define the *lower residuals* as *E_i_* = *q̂_α/_*_2_(*Xi*)−*Y_i_* and *upper residuals R_i_* = *Y_i_*−*q̂*_1_*_−α/_*_2_(*X_i_*) where *i* ∈ *N_C_*. Denote the size of the calibration set as *n_C_* where *n_C_* = |*N_C_*|. We estimate calibration constants *ĉ_α/_*_2_ as the (1 − *α/*2)(1 + 1*/n_C_*)-th empirical quantile of the *E_i_* and *ĉ*_1_*_−α/_*_2_ and (1 − *α/*2)(1 + 1*/n_C_*)-th empirical quantile of the *R_i_*, *i* ∈ *N_C_*. These empirical quantiles are computed as the ⌈(1−*α/*2)(*n_C_* +1)⌉ smallest of *E_i_* and *R_i_* respectively, *i* ∈ *N_C_*. Under exchangeability of the full data and new observation and continuous-valued residuals, Romano et al. (2019b) prove that the reference interval *Ĉ*(*X*) = [*q̂_α/_*_2_(*X*) − *ĉ_α/_*_2_*, q̂*_1_*_−α/_*_2_(*X*) + *ĉ*_1_*_−α/_*_2_] has the properties

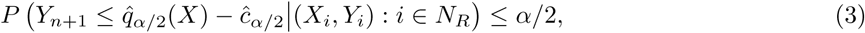

and

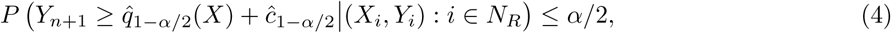

where the probability is taken over the calibration set and the new observation. That is, the reference interval *Ĉ*(*X*) satisfies Properties 1 and 2 and is thus a valid reference interval.

In our setting however, the reference set and calibration set may have different distributions. In neuroimaging, this situation occurs when a reference interval is fit in reference data and applied to a new dataset, which differs in study design, MRI acquisition, or image preprocessing. Now, we assume that the reference and calibration sets differ in distribution such that (*X_i_, Y_i_*) ∼ *P* , *i* ∈ *N_R_* and (*X_i_, Y_i_*) ∼ *Q*, *i* ∈ *N_C_*. This setting violates the original assumptions of both conformal prediction and CQR since the full data is no longer exchangeable (Lei et al., 2018; Romano et al., 2019b). However, the proof of Equations 3 and 4 only depends on exchangability of the calibration set and new observation.

Here, we assume the calibration set (*X_i_, Y_i_*), *i* ∈ *N_C_* and new observation (*X_n_*_+1_*, Y_n_*_+1_) are exchangeable and distributed as *Q*. We first use the reference set (*X_i_, Y_i_*)*, i* ∈ *N_R_* to estimate a naïve reference interval *C*(*X*) = [*q̂_α/_*_2_(*X*)*, q̂*_1_*_−α/_*_2_(*X*)]. Next, we aim to *calibrate C*(*X*) to obtain a valid reference interval for the new observation (*X_n_*_+1_*, Y_n_*_+1_). Using the estimation procedure from CQR, we estimate calibration constants *ĉ_α/_*_2_ as the (1 − *α*)(1 + 1*/n_C_*)-th empirical quantile of the *E_i_* and *ĉ*_1_*_−α/_*_2_ and (1 − *α*)(1 + 1*/n_C_*)-th empirical quantile of the *R_i_*, *i* ∈ *N_C_*. We call this method Reference interval calibration via conFormal prediction (ReForm) and our derived interval a *ReFormed* reference interval given by

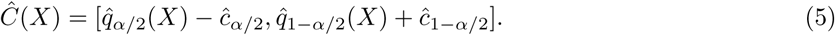

Using the same definition of lower residuals *E_i_* and upper residuals *R_i_* from CQR, we know that *E_i_* and *R_i_* are both continuous-valued and exchangeable among *i* ∈ *N_C_* and *i* = *n* + 1. Thus, from Romano et al. (2019b), our ReFormed reference interval satisfies Equations 3 and 4 where the probability is taken over the calibration set and the new observation. Therefore, our Reformed reference interval is a valid reference interval satisfying Properties 1 and 2. Note that first, we make no assumptions on the reference set distribution *P* or the calibration set distribution *Q*. Second, if the naïve reference interval *C*(*X*) is available, the reference set is not required to obtain the ReFormed reference interval.

We implement our method in the *ReForm* R package available via GitHub (https://github.com/andy1764/ReForm), which includes functions tailored to widely-used regression models.

### 2.3 Alternative methods

Several alternative methods have been proposed in the literature for either adjusting data or adjusting models in order to mitigate differences between datasets. Here, we compare ReForm to methods from three categories: refitting, harmonization, and model adjustment.

*Refitting* is the simplest solution where, instead of using the reference interval, we fit new intervals entirely in the calibration set. Refitting is recommended by the CLSI when a reference interval fails to be verified in a new sample (Clinical and Laboratory Standards Institute, 2008). Here, this corresponds to a reference interval 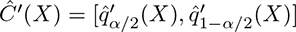 where 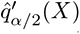 and 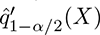 are estimated conditional quantiles fit in the calibration set, *i* ∈ *N_C_*. Refitting is not guaranteed to yield a valid reference interval and may provide inaccurate fits particularly in small calibration sets.

*Harmonization* methods aim to transform the new observation to have the same conditional distribution as the reference data. In our setting where the reference set (*X_i_, Y_i_*) ∼ *P* = *P_X_P_Y |X_* , *i* ∈ *N_R_*, this corresponds to estimating a function *h* such that *h*(*Y_n_*_+1_*, X_n_*_+1_) ∼ *P_Y |X_* . For harmonization, we can obtain an estimate *ĥ* from the full data *i* = 1, 2*, . . . , n* and apply to the (*n*+1)-th observation. The form of *ĥ* depends on the chosen harmonization method, which can employ: linear models (Johnson et al., 2007); generalized additive models (Pomponio et al., 2020); generalized additive models for location, scale, and shape (GAMLSS; Gardner et al., 2025); deep learning (Hu et al., 2024); and other models (Beer et al., 2020; Hu et al., 2023; Ling et al., 2022). Recent work suggests that fitted harmonization parameters from ComBat and several of its extensions can be applied to new sites without using the reference set (Xin et al., 2025).

*Model adjustment* methods fit a reference interval in the full data and apply that interval directly to the new observation. The specific model used to fit the interval varies, but typically includes a term for *batch*. In neuroimaging, batch is typically used to indicate the study site or MRI scanner for each observation.

Assuming that the new observation is from a batch in the full data, we can directly estimate *C*(*X*) as described previously and apply to the new observation (*X_n_*_+1_*, Y_n_*_+1_), where here *X_i_, i* = 1, 2*, . . . , n* + 1 includes indicators for batch. In a recent study, Bethlehem et al. (2022) develop a new method based on GAMLSS with a random effect for batch to fit the random effect in a new batch, without access to the reference set. However, model adjustment methods do not guarantee validity of reference intervals.

Figure 1 illustrates unadjusted reference intervals (Unadjusted, 1a), harmonization via ComBatLS (1b), refitting in the calibration set (Refit, 1c), and ReForm (1d), all using empirical data from our study.

**Figure 1:**
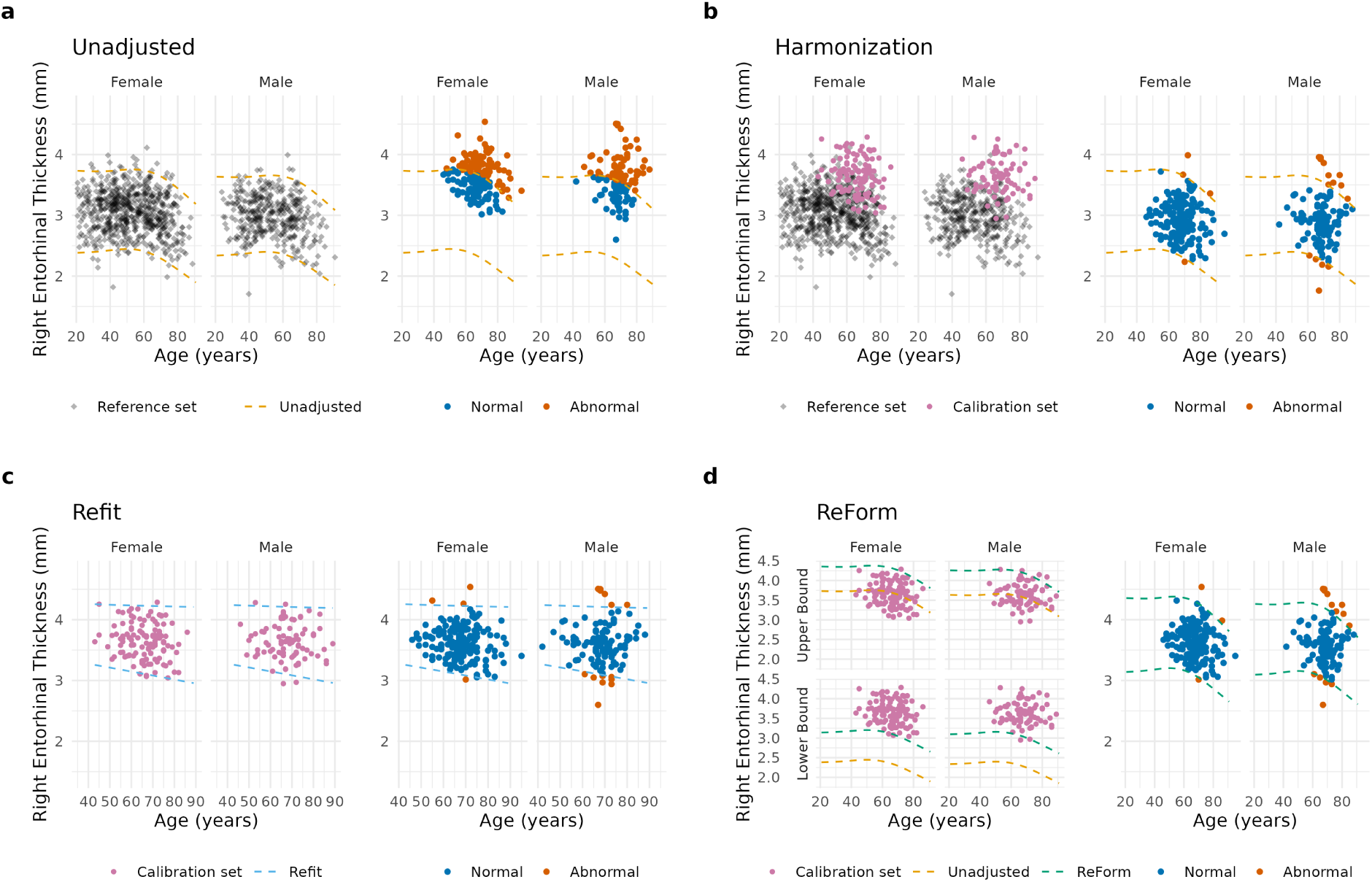
Illustration of methods employed in Experiment 1, which fitted reference intervals in ARWiBO 1.0 T and applied them to OASIS 3.0 T scans. All reference intervals were applied to the same validation set consisting of 288 OASIS 3.0 T scans, represented as colored points where blue is inside the reference interval (normal) and orange is outside (abnormal). Data shown is from a single run from Experiment 1. (a) 1032 ARWiBO scans (diamond points) were used to fit 95% reference intervals using GAMLSS (left, light orange dashed line), which were directly applied to OASIS 3.0 T scans yielding 152 (52.7%) outside the interval. (b) 1032 ARWiBo scans were combined with 200 OASIS 3.0 T scans (pink) to harmonize across sites (left), which shifted the validation set (right) yielding 19 outside (6.5%) (c) A 95% reference interval was fit in 200 OASIS 3.0 T scans (pink) using GAMLSS (left, light blue dashed line) and applied to the validation set yielding 18 outside (6.3%). (d) The upper and lower bounds of the reference interval from (a) were calibrated using Reference interval calibration via conFormal prediction (ReForm, left, green dashed line) in 200 OASIS 3.0 T scans and applied to the validation set yielding 17 outside (5.9%).

### 2.4 Considerations for ReForm

ReForm only requires exchangeability of the calibration set and new observation to yield valid reference intervals. However, ReForm does not guarantee that the ReFormed reference interval accurately reflects the underlying distribution of the new observation. That is, for a new observation (*X_n_*_+1_*, Y_n_*_+1_) ∼ *Q* where *Q* = *Q_X_Q_Y |X_* , the ReFormed interval *Ĉ*(*X*) from Equation 5 is not guaranteed to be close to the actual population *α/*2 or 1 − *α/*2 quantiles of *Q_Y |X_* . ReForm adjusts the upper bound and lower bound of the naïve reference interval *C*(*X*) by separate constants. This adjustment would be ideal if the *α/*2 or 1 − *α/*2 quantiles of the conditional distributions of the reference set *P_Y |X_* , where *P* = *P_X_P_Y |X_* , and calibration set *Q_Y |X_* differ by separate constants. If that relationship is true and the naïve reference interval is close to the true *α/*2 or 1 − *α/*2 quantiles of *P_Y |X_* , then the ReFormed reference interval should be close to the quantiles of *Q_Y |X_* . This situation is similar to a common assumption in harmonization methods, which is that batches differ in mean and variance (Beer et al., 2020; Gardner et al., 2025; Johnson et al., 2007; Pomponio et al., 2020). This situation is also comparable to the assumptions of ConQuR, which adjusts the conditional quantiles across batches to perform harmonization (Ling et al., 2022), and GAMLSS with random effects, which are typically specified for both the mean and variance (Bethlehem et al., 2022). In our ReForm R package, we provide diagnostic plots based on the upper and lower residuals for qualitative evaluation.

Since our primary goal is to provide a reference interval calibration method that controls FPR in a new individual, we do not evaluate our method based on recovery of the true quantiles of *Q_Y |X_* . However, we compare ReForm to refitting (Refit) throughout all experiments, where refitting uses the calibration set to obtain an estimate of the population quantiles of *Q_Y |X_* .

Our setting is a special case of covariate shift and distribution shift, where the new observation is drawn from a different distribution than the full data (Barber et al., 2023; Tibshirani et al., 2019). In our setting however, we have a calibration set available that is drawn from the same distribution as the new observation. Therefore, the results of CQR hold when using that calibration set. However, distribution shift relative to the calibration set could occur in situations where the sampling scheme or data acquisition changes over the course of an MRI study.

### 2.5 Image acquisition and preprocessing

All data in this study are from the Lifespan Brain Chart Consortium (LBCC) with preprocessing details previously published (Bethlehem et al., 2022; Dorfschmidt et al., 2025). In summary, all studies collected T1-weighted images and certain studies collected T2-FLAIR images as well. When both T1-weighted and T2-FLAIR images were available, FreeSurfer’s combined T1-T2 *recon-all* was used to extract volumetric and cortical measures. When only T1-weighted images were available, FreeSurfer’s standard *recon-all* was used instead. Briefly, *recon-all* involved non-uniformity correction, projection to Talairach space, intensity normalisation, skull-stripping, automatic tissue and subcortical segmentation, surface interpolation, tessellation and registration. Different FreeSurfer versions were used across the LBCC studies with additional details available elsewhere (Bethlehem et al., 2022; Dorfschmidt et al., 2025). All data analyzed in this work used FreeSurfer version 6.0.142.

Subject details were provided by the individual studies included in the LBCC. We used a subset of baseline scans from AD studies for our experiments. Our inclusion and study-specific diagnostic criteria are detailed under the descriptions of each experiment in this study.

### 2.6 Resampling experiments in LBCC data

We chose to use right entorhinal cortical thickness as our primary outcome. Our rationale is that atrophy in this region has been identified in mild cognitive impairment preceding AD with further reductions across stages of AD (Juottonen et al., 1999; Tapiola et al., 2008). We chose to examine cortical thickness owing to its high sensitivity to MRI acquisition and preprocessing differences (Chen et al., 2022; Fortin et al., 2018; Han et al., 2006).

Across all experiments, we evaluated methods via FPR. For a valid reference interval, FPR should be controlled at *α* across randomly sampled new observations for a given calibration set. However, empirical FPR may vary depending on the demographics and outcomes included in the calibration set. FPR should also depend on the size of the calibration set, following results from conformal prediction that size impacts the upper bound on coverage (Lei et al., 2018; Romano et al., 2019b). Furthermore, FPR is computed from the data, so the theoretical guarantees of ReForm may not correspond to exact control of FPR. For comparison against the target FPR, we applied one-sample Wilcoxon signed-rank tests. For comparison between methods, we used Wilcoxon signed-rank tests across pairs of methods applied to the same data splits. All tests were performed with a significance level of 0.05.

Our experiments estimated 90%, 95%, and 99% reference intervals which have target FPRs of 0.1, 0.05, and 0.01 respectively. In all experiments, we designated separate reference dataset and test datasets as different sites from the LBCC data. Then, we randomly drew reference sets from the reference dataset and calibration sets from the test dataset, both without replacement. Across experiments, reference set sizes varied across {50, 200*, n_T_* } where *n_T_* refers to the size of the entire reference dataset. Calibration sets varied across {20, 40, 80, 120, 160, 200, 300, 400} throughout all experiments. Once the calibration set was drawn, the validation set consisted of all remaining observations in the test dataset. We primarily used GAMLSS to fit reference intervals assuming a Box-Cox t-distribution (BCT) with a fixed effect for biological sex and a smooth age term with default settings in both the mean and variance. The reference interval from GAMLSS was estimated from the population quantiles of the estimated BCT distributions. In Supplementary Materials, we report results using additive quantile regression instead, including a fixed effect for biological sex and a smooth age term with default settings. Each experiment was repeated 1,000 times for each unique combination of reference set size, calibration set size, and model choice.

We compared ReForm to: the unadjusted naïve reference interval (Unadjusted), refitting reference intervals in the calibration set (Refit); two harmonization methods (ComBat-GAM and ComBatLS); and GAMLSS-based model adjustment where appropriate. ComBat-GAM and ComBatLS were chosen since both account for nonlinear age effects, but ComBatLS additionally models potential covariate effects in the scale of measurements (Gardner et al., 2025). GAMLSS-based model adjustment only applied to settings where the reference interval was constructed using a GAMLSS model and the reference dataset had multiple sites. For model adjustment, our chosen GAMLSS model (GAMLSS-RE) assumed a BCT distribution and specified fixed effects for biological sex, a smooth age term with default settings, and a random effect for batch, all in both the mean and variance. Both the harmonization and model adjustment methods employed can be fit without the reference set; however, omitting the reference introduces error (Bethlehem et al., 2022; Xin et al., 2025). Here, we compared ReForm to methods that use the reference set in order to assess performance against the optimal implementation of existing methods.

GAMLSS was implemented via the *gamlss2* R package v0.1-0 (https://github.com/gamlss-dev/gamlss2). Additive quantile regression was fit using the *qgam* R package v2.0.0 on CRAN (Fasiolo et al., 2021). ComBat-GAM and ComBatLS were both implemented using the *CombatFamily* R package v0.2.1 (https://github.com/andy1764/ComBatFamily), which allows for estimated harmonization parameters to be applied to the validation set. ComBat-GAM was fit using a smooth term for age and fixed effect for sex, both specified using the *gam* function from the *mgcv* R package v.1.9-1. ComBatLS was fit using smooth terms for age (*ps* function) and fixed effects for sex in both the mean and variance using the *gamlss* R package v.5.4-22. Since our outcome involved a single outcome, ComBat-GAM and ComBatLS are implemented without the empirical Bayes shrinkage across outcomes (Johnson et al., 2007).

### 2.7 Experiment 1: Different study, different acquisition

Our first experiment used 1.0 T scans from the Alzheimer’s Disease Repository Without Borders (ARWiBo; Frisoni et al., 2009) study to fit reference intervals and applied them to 3.0 T scans from the Open Access Series of Imaging Studies (OASIS 3.0 T; LaMontagne et al., 2019; Marcus et al., 2007, 2010). Our goal for this experiment was to assess how severely FPR could be impacted by study differences, including a large magnetic field strength difference (1.0 T vs 3.0 T) in MRI acquisition. The ARWiBo study defined cognitively normal (CN) as the absence of subjective memory complaints, objective memory impairment based on several standard exams, and AD assessed via the NINCDS-ADRDA criteria Frisoni et al. (2009); McKhann et al. (1984). The OASIS study defined CN as subjects assessed via the Clinical Dementia Rating (CDR) scale (LaMontagne et al., 2019; Morris, 1993) to have a CDR of less than 0.5. Here, the aforementioned GAMLSS model adjustment method could not be applied since the reference dataset only contains a single batch.

### 2.8 Experiment 2: Same study, different acquisition

Our second experiment used 1.5 T scans from OASIS study (OASIS 1.5 T; Marcus et al., 2007, 2010) to fit reference intervals and applied them to the OASIS 3.0 T. Unlike our first experiment, the reference and test dataset here were both from the OASIS study with identical inclusion criteria, the same definition of CN, and comparable acquisition parameters (LaMontagne et al., 2019). However, we anticipated that there would still notable differences due to the difference in magnetic field strength (1.5 T vs 3.0 T). In this experiment, we compared ReForm to refitting (Refit) and ComBatLS. Again, the GAMLSS model adjustment method could not be used in this setting.

### 2.9 Experiment 3: Multi-site reference dataset

Our third experiment used multi-site reference data from the National Alzheimer’s Coordinating Center (NACC), which used several 3.0 T scanners across multiple study sites with details reported previously (Morris et al., 2006). We fitted reference intervals in NACC and applied these reference intervals to the OASIS 3.0 T. The NACC database used its own diagnostic criteria based on several screening tools, including the CDR, to categorize subjects as either CN, mild cognitive impairment, or AD patients (Morris et al., 2006). In this study, we included CN from both studies and AD subjects from the OASIS 3.0 T, where the OASIS study defined AD as subjects having a CDR of 0.5 or greater (LaMontagne et al., 2019).

Here, we first compared in CN subjects the following methods: ReForm, the unadjusted naïve reference interval (Unadjusted), refitting (Refit), the same two harmonization methods (ComBat-GAM and ComBatLS), and GAMLSS model adjustment treating batch as a random effect (GAMLSS-RE). Batches included the 22 NACC study sites and OASIS 3.0 T. Then, we drew calibration sets from OASIS 3.0T CN subjects and compared all methods in the OASIS 3.0 T AD patients. Our goal was to measure the *positive rate* (PR) estimated as the proportion of OASIS 3.0 T AD patients with observations outside a given reference interval. We aimed to demonstrate similar or increased PR while controlling FPR at our designated level. As a supplementary analysis, we also examine PR subsetting to trials where the FPR close to our target FPR (*α*), within *α* ± 0.2 × *α*. Finally, we applied our NACC reference intervals to the ARWiBo 1.0 T study described previously.

## 3 Results

### 3.1 Data subset from the LBCC

Table 1 reports the sample used across all experiments labeled by the experiment (e.g. Exp. 1 Reference refers to Experiment 1 reference set). For this investigation, we only included subjects that are labeled as either cognitively normal (CN) or Alzheimer’s disease (AD) based on the individual study’s criteria detailed in Section 2. Experiment 1 included CN subjects from the ARWiBo and OASIS 3.0 T studies (*n* = 1520). Experiment 2 included CN subjects from the OASIS 1.5 T and OASIS 3.0 T studies (*n* = 588). Experiment 3 included CN subjects from the NACC and OASIS 3.0 T studies (*n* = 3311 across 23 study sites) and AD subjects from the OASIS 3.0 T study (*n* = 199).

**Table 1:** Subject characteristics by study and site.

|  | ARWiBo | OASIS |  | NACC |
| --- | --- | --- | --- | --- |
|  | 1.0 T<br>(Exp. 1 Reference) | 1.5 T<br>(Exp. 2 Reference) | 3.0 T<br>(Exp. 1-3 Test) | All 22 Sites<br>(Exp. 3 Reference) |
| **Sample size** | 1155 | 120 | 687 | 4530 |
| **Age (years)** |  |  |  |  |
| Mean (SD) | 55.0 (16.1) | 65.9 (11.3) | 69.8 (8.92) | 71.8 (10.6) |
| Median [Min, Max] | 54.8 [20.8, 91.8] | 66.0 [45.0, 95.0] | 70.0 [42.0, 95.0] | 72.5 [19.7, 101] |
| **Sex** |  |  |  |  |
| Female | 735 (63.6%) | 83 (69.2%) | 383 (55.7%) | 2800 (61.8%) |
| Male | 420 (36.4%) | 37 (30.8%) | 304 (44.3%) | 1730 (38.2%) |
| **Diagnosis** |  |  |  |  |
| CN | 1032 (89.4%) | 100 (83.3%) | 488 (71.0%) | 2823 (62.3%) |
| AD | 123 (10.6%) | 20 (16.7%) | 199 (29.0%) | 1707 (37.7%) |

Abbreviations: Exp., Experiment; ARWiBo, Alzheimer’s Disease Repository Without Borders; OASIS, Open Access Series of Imaging Studies; NACC, National Alzheimer’s Coordinating Center; SD, Standard deviation; CN, Cognitively normal; AD, Alzheimer’s disease

### 3.2 ReForm versus refitting intervals

In general, we found that ReForm outperformed the Unadjusted and Refit methods for controlling FPR. Figure 2 shows FPR as medians and interquartile ranges (IQR) from Experiment 1. For 95% reference intervals with the full reference dataset and calibration set size of 40, the Unadjusted method resulted in median FPR of 0.538, far exceeding the target FPR of 0.05. ReForm demonstrated better control of FPR (median FPR 0.042, IQR 0.022-0.065) than Refit (median FPR 0.104, IQR 0.074-0.147; Refit vs. Reform Wilcoxon signed-rank test *p <* 0.001). In a larger calibration set of size 400, ReFit (median FPR 0.057, IQR 0.034-0.068) and ReForm (median FPR 0.045, IQR 0.034-0.068) showed more comparable performance. For 99% reference intervals and a small calibration set of size 40, both Refit (median FPR 0.040, IQR 0.022-0.069) and ReForm (median FPR 0.040, IQR 0.022-0.067) failed to control FPR at *α* = 0.01. With 160 calibration samples, Refit still showed poor control (median FPR 0.015, IQR 0.009-0.021) while ReForm significantly outperformed (median FPR 0.009, IQR 0.006-0.018; Refit vs ReForm *p <* 0.001). ReForm was uniquely able to achieve FPR across trials not significantly different from 0.01 across all reference set sizes at a calibration set size of 400. Using additive quantile regression instead of GAMLSS, we found that ReForm controlled the FPR of 95% and 90% reference intervals better than Refit for small calibration sets and was not significantly different from the target FPR in large calibration sets (Supplementary Figure 1).

**Figure 2:**
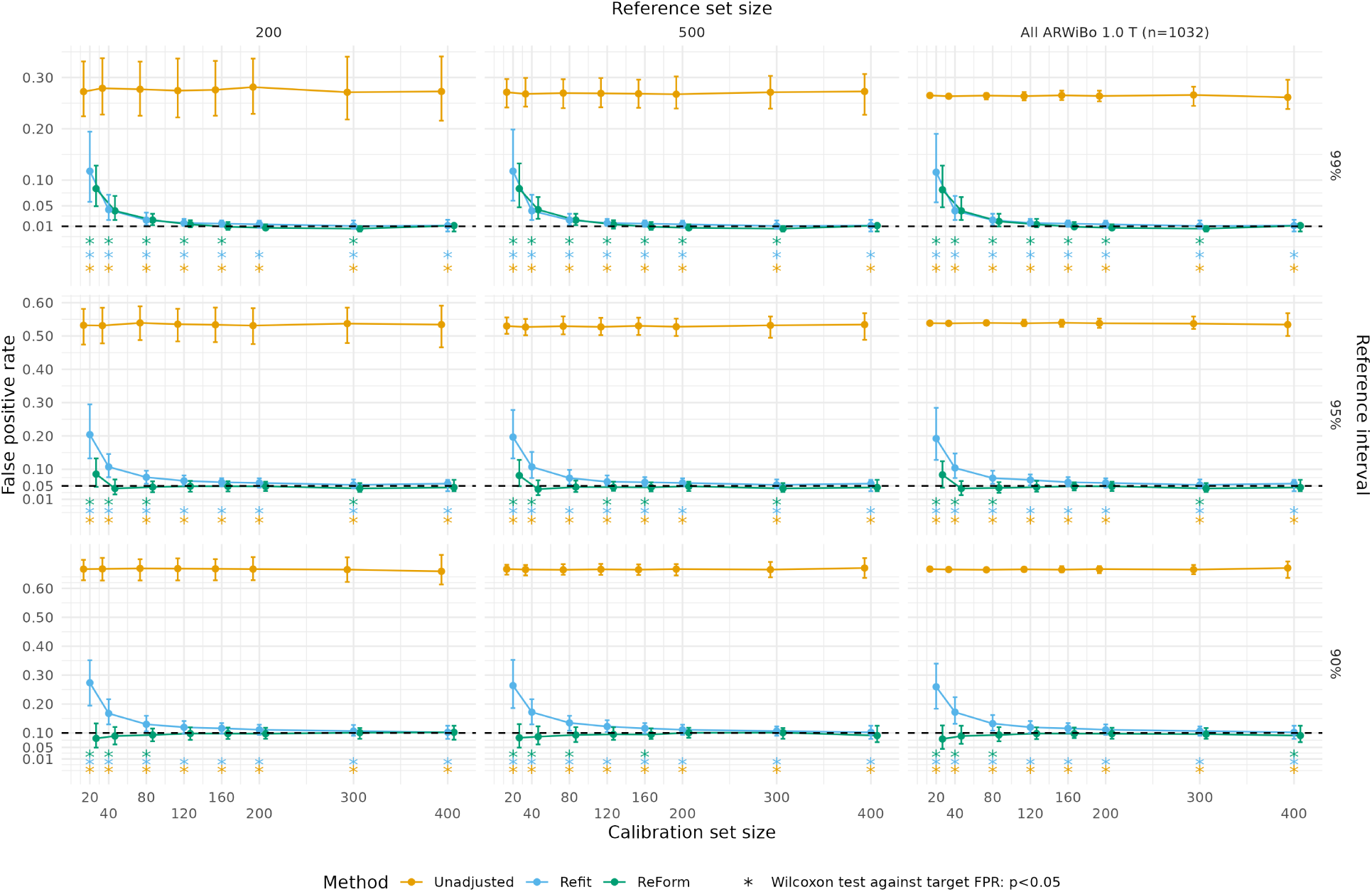
False positive rates across unadjusted, refit, and ReFormed reference intervals from Experiment 1, where we fitted reference intervals in ARWiBO 1.0 T scans and applied them to OASIS 3.0 T scans. Median and interquartile range are plotted across 1,000 trials. Plots assess the validity of estimated 99% reference intervals (top panel, target FPR of 0.01); 95% reference intervals (middle panel, target FPR of 0.05; and 90% reference interval (bottom panel, target FPR of 0.1). Asterisks used to show significant differences from the target FPR (dashed line) using a one-sample Wilcoxon signed-rank test. Unadjusted represents direct application of reference intervals fit using GAMLSS. Refit refers to refitting using GAMLSS in the calibration set of a designated size. ReForm is our proposed method which calibrates the unadjusted reference interval. All FPR values are computed in the same OASIS 3.0 T scans with varying sample size left after holding out the calibration set. In Refit, some trials yielded errors during fitting and are not shown.

### 3.3 ReForm versus statistical harmonization

We first observed that ComBatLS notably outperformed ComBat-GAM for FPR control, but both harmonization methods failed to control FPR in certain settings. Figure 3 shows FPR of ComBat-GAM, ComBatLS, and ReForm across trials in Experiment 1, where intervals are fit in the ARWiBo 1.0 T and applied to OASIS 3.0 T scans. For 90% reference intervals, ComBat-GAM exhibited dependence on the reference set size. Fixing the calibration set size at 200 and comparing reference set sizes of 200 and 1033, ComBat-GAM median FPR goes from 0.128 (IQR 0.104-0.156) to 0.104 (IQR 0.087-0.122). Comparing ComBat-GAM and ComBatLS in an ideal setting where both methods have access to 500 reference samples and a large calibration set of size 400, ComBatLS exhibited significantly lower FPR (median FPR 0.102, IQR 0.068-0.125) than ComBat-GAM (median FPR 0.114, IQR 0.091-0.148; one-sided Wilcoxon signed-rank test *p <* 0.001). Comparing ComBatLS versus ReForm for 90% reference intervals, ComBatLS demonstrated poor control of FPR in calibration sets of size 120 (median FPR 0.120, IQR 0.101-0.139) while ReForm exhibited FPR not significantly different than the target FPR of 0.1 (median FPR 0.098, IQR 0.079-0.120; one-sample Wilcoxon signed-rank test against 0.1, *p* = 0.48).

**Figure 3:**
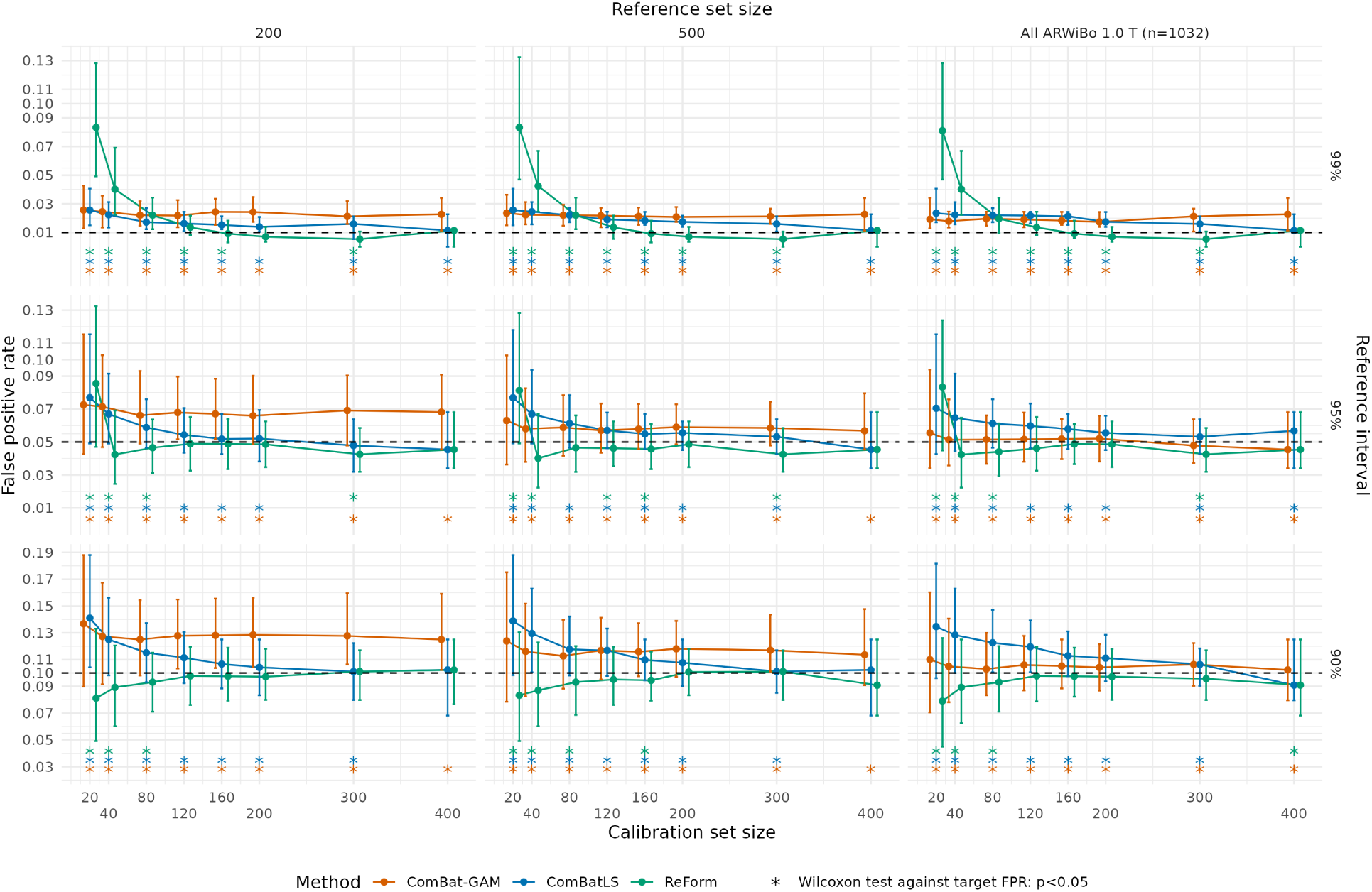
False positive rates across harmonization methods and ReForm from Experiment 1, where we fitted reference intervals in ARWiBO 1.0 T scans and applied them to OASIS 3.0 T scans. Median and interquartile range are plotted across 1,000 trials. Plots assess the validity of estimated 99% reference intervals (top panel, target FPR of 0.01); 95% reference intervals (middle panel, target FPR of 0.05; and 90% reference interval (bottom panel, target FPR of 0.1). Asterisks used to show significant differences from the target FPR (dashed line) using a one-sample Wilcoxon signed-rank test. ComBat-GAM and ComBatLS are fit on the entire reference set combined with the calibration set. ReForm is our proposed method which calibrates the unadjusted reference interval. All FPR values are computed in the same OASIS 3.0 T scans with varying sample size left after holding out the calibration set.

The superior performance of ReForm over ComBatLS was more stark in Experiment 2, which fits reference intervals in the OASIS 1.5 T for application to OASIS 3.0 T. As hypothesized, we observed that the FPRs of unadjusted intervals in Experiment 2 were closer to the target FPRs compared to Experiment 1 (Figure 3 vs. Figure 4). However, we found that ComBatLS showed poor FPR control even in ideal settings. For 95% reference intervals with access to the full reference set and calibration set size of 400, ComBatLS exhibits FPR significantly different than the target FPR of 0.05 (median 0.080, IQR 0.057-0.102; one-sample Wilcoxon signed-rank test against 0.05, *p <* 0.001). In this setting, ReForm achieves FPR close to *α* = 0.05 (median FPR 0.045, IQR 0.034-0.068; *p* = 0.88). For 99% reference intervals in the same setting, performance of ComBatLS (median FPR 0.034, IQR 0.023-0.045) was close to Unadjusted (median FPR 0.034, IQR 0.023-0.045). In contrast, ReForm demonstrated FPR not significantly different than the target FPR of 0.01 (median FPR 0.011, IQR 0.000-0.011; one-sample Wilcoxon signed-rank test against 0.01, *p* = 0.74). Moreover, we observed that ComBatLS depended on the reference set size while ReForm did not. Moving from 50 reference samples to the full reference set to fit 95% reference intervals, ComBatLS median FPR decreases substantially from 0.102 (IQR 0.068-0.136) to 0.080 (IQR 0.057-0.102) whereas ReForm is stable at 0.045 (IQR 0.034-0.068) for both reference set sizes with calibration set size of 400.

**Figure 4:**
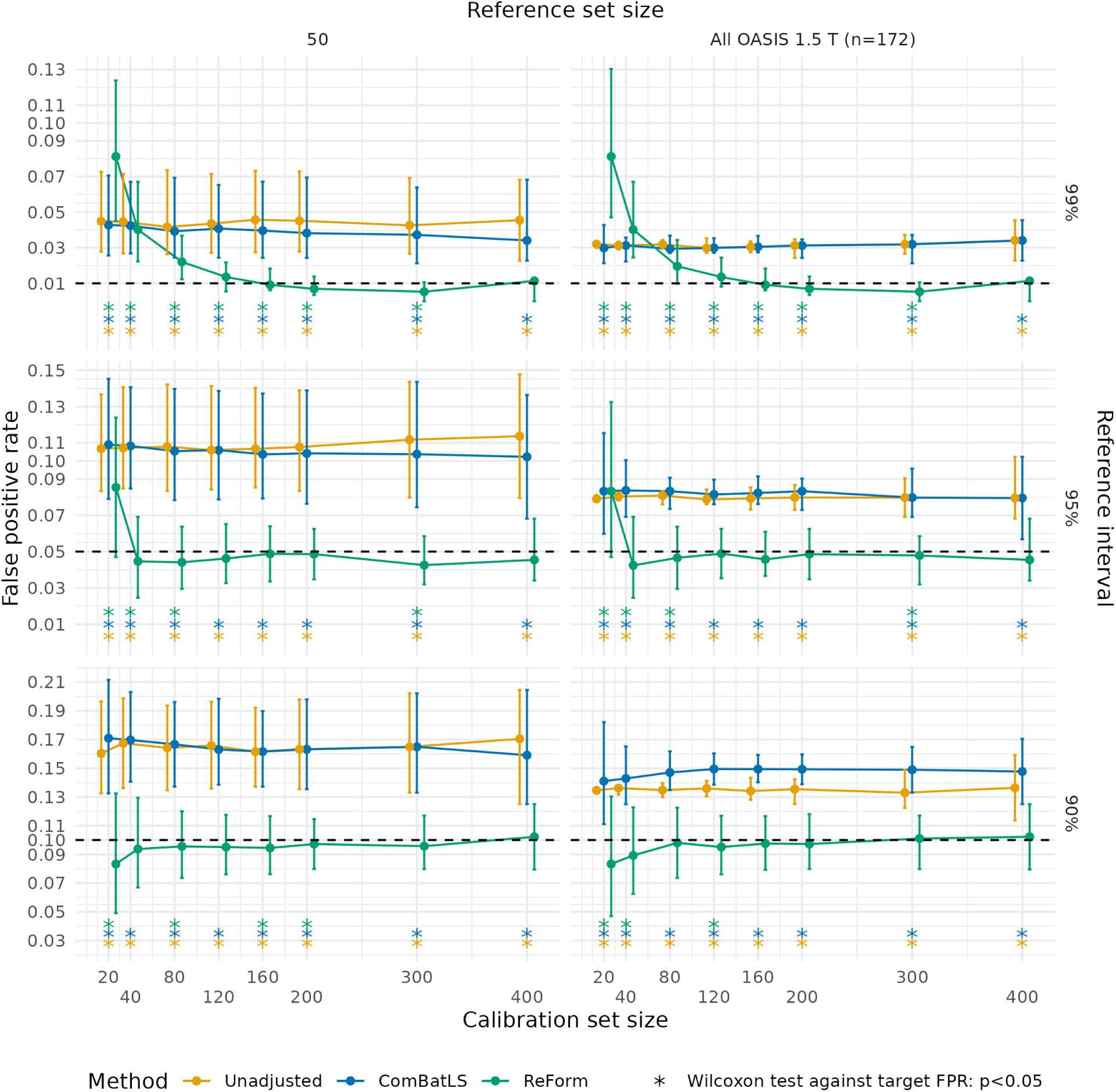
False positive rates across unadjusted, ComBatLS, and ReForm for fitting reference intervals from Experiment 2, where we fitted reference intervals in OASIS 1.5 T scans and applied them to OASIS 3.0 T scans. Median and interquartile range are plotted across 1,000 trials. Plots assess the validity of estimated 99% reference intervals (top panel, target FPR of 0.01); 95% reference intervals (middle panel, target FPR of 0.05; and 90% reference interval (bottom panel, target FPR of 0.1). Asterisks used to show significant differences from the target FPR (dashed line) using a one-sample Wilcoxon signed-rank test. Unadjusted represents direct application of reference intervals fit using GAMLSS. ComBatLS is fit on the entire reference set combined with the calibration set. ReForm is our proposed method which calibrates the unadjusted reference interval. All FPR values are computed in the same OASIS 3.0 T scans with varying sample size left after holding out the calibration set.

In Experiment 3, Figure 5 shows that ComBatLS fit across NACC study sites and OASIS 3.0 T failed to control FPR across most settings. For instance, ComBatLS for 95% reference intervals showed poor control in both small calibration sets of size 40 (median FPR 0.067, IQR 0.049-0.089; one-sample Wilcoxon signed-rank test against 0.05, *p <* 0.001) and large calibration sets of size 400 (median FPR 0.057, IQR 0.045-0.068; *p <* 0.001).

**Figure 5:**
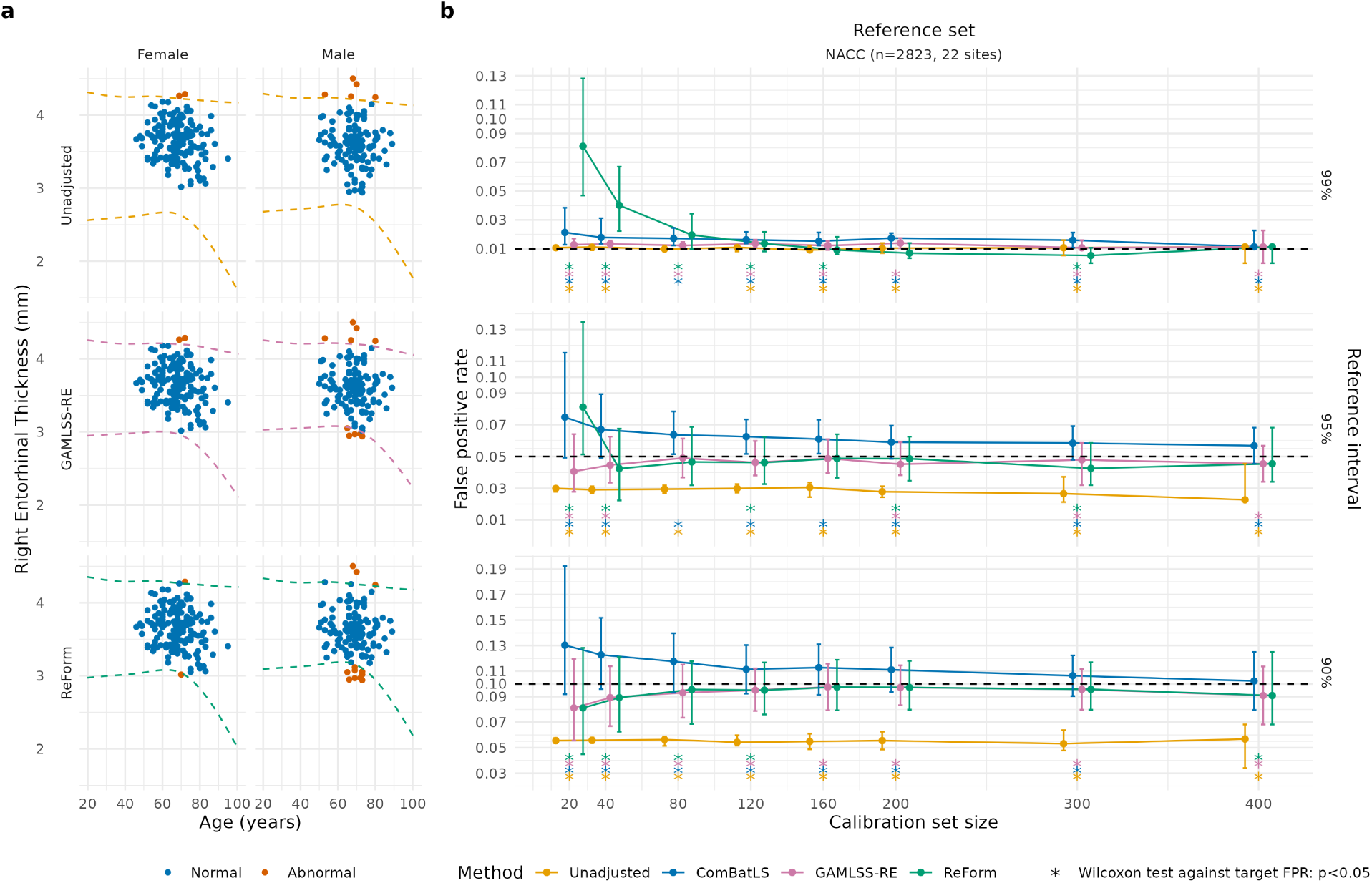
Application of reference intervals fitted in the NACC reference dataset to OASIS 3.0 T healthy subjects in Experiment 3. (a) Illustration of methods using a single trial from Experiment 3, where we fitted reference intervals in the multi-site NACC study and applied them to OASIS 3.0 T scans. 288 OASIS 3.0 T healthy scans are shown. Direct application of 95% reference intervals (Unadjusted) yielded 7 outside (2.4%). A 95% reference interval constructed using GAMLSS with a random effect across the 2928 NACC scans and 200 OASIS 3.0 T healthy scans yielded 12 outside (4.2%). A ReFormed reference interval yielded 14 outside (4.9%) (b) False positive rates across unadjusted, ComBatLS, GAMLSS-RE and ReForm for Experiment 3. Median and interquartile range are plotted across 1,000 trials. Plots assess the validity of estimated 99% reference intervals (top panel, target FPR of 0.01); 95% reference intervals (middle panel, target FPR of 0.05; and 90% reference interval (bottom panel, target FPR of 0.1). Asterisks used to show significant differences from the target FPR (dashed line) using a one-sample Wilcoxon signed-rank test. Unadjusted represents direct application of reference intervals fit using GAMLSS. ComBatLS is fit on the entire reference set combined with the calibration set. GAMLSS-RE refers to fitting a GAMLSS with random effect for study site across the combined reference and calibration sets. ReForm is our proposed method which calibrates the unadjusted reference interval. All FPR values are computed in the same OASIS 3.0 T scans with varying sample size left after holding out the calibration set.

### 3.4 ReForm versus GAMLSS in pooled multi-site data

Compared to fitting GAMLSS across all sites with a random effect for site (GAMLSS-RE), we found that ReForm showed less stable FPR control in small calibration sets but more precise control in larger calibration sets. Figure 5a illustrates these methods in a single trial from Experiment 3. Figure 5b compares FPR across Unadjusted, ComBatLS, and fitting GAMLSS across all sites with a random effect for site (GAMLSS-RE), and ReForm. In Unadjusted, we observed in 99% reference intervals that the FPR is already close to the target of 0.01 (Unadjusted median FPR 0.011). GAMLSS-RE performed consistently even across small calibration samples taken from OASIS 3.0 T. For example, 99% reference intervals constructed via GAMLSS-RE controlled FPR consistently even with 20 calibration samples (median FPR 0.013, IQR 0.011-0.017), whereas ReForm did not (median FPR 0.081, IQR 0.047-0.128, Wilcoxon rank-sum test *p <* 0.001). We found that ReForm was able to achieve FPR control not significantly different from the target FPRs for 95% and and 99% reference intervals with 400 calibration samples, while GAMLSS-RE did not (Figure 5). ReForm was not significantly different than the target FPR for 160 calibration samples in 90% and 95% reference intervals (Figure 5). In Supplementary Materials, we observe similar results in application to the ARWiBo study where ReForm achieves comparable performance to GAMLSS-RE at 80 samples for 90% reference intervals, 120 for 95% reference intervals, and 160 samples for 99% reference intervals (Supplementary Figure 3a).

### 3.5 Reference intervals applied to Alzheimer’s disease patients

In applying reference intervals to AD patients, ReForm and GAMLSS-RE both considerably outperformed Unadjusted in terms of positive rate (PR), the proportion of AD patients with right entorhinal thickness outside of a given reference interval. Figure 6a illustrates a ReFormed reference interval compared to GAMLSS-RE and direct application of unadjusted intervals (Unadjusted), all from a single run of Experiment 3. Figure 6b compares PR across Unadjusted, GAMLSS-RE, and ReForm across all trials. Results for *α* = 0.01 are omitted since we observed that neither method controlled FPR adequately; less than 20% of trials had both methods yield FPR between 0.008 and 0.012 (Supplementary Figure 4a). First, we observed that the unadjusted reference intervals yield a median PR below 0.15 across 90% (median FPR 0.141), 95% (median FPR 0.085), and 99% reference intervals (median FPR 0.020). Next, we compared GAMLSS-RE and ReForm at a large calibration set size of 200, where both methods showed good control of FPR. Comparing GAMLSS-RE and ReForm for 95% intervals, both showed similar performance (GAMLSS-RE median PR 0.201, IQR 0.196-0.201; ReForm median PR 0.206, IQR 0.186-0.221). For 90% reference intervals, ReForm achieved a higher median PR (0.296, IQR 0.251-0.312) than GAMLSS-RE (0.256, IQR 0.251-0.271; one-sided Wilcoxon signed-rank test *p <* 0.001). This PR difference corresponded to median 5 (IQR 1-9) more AD subjects being labelled as abnormal based on the ReFormed reference interval versus the GAMLSS-RE interval. Sup-plementary Figure 4b shows performance across trials where FPR for both methods is controlled within 20% of the target FPR, where we observed similar results. Supplementary Figures 3b and 5 show results for PR when applying reference intervals instead to the ARWiBo AD patients (*n* = 123), generally showing similar results despite the severely inflated PR in Unadjusted intervals.

**Figure 6:**
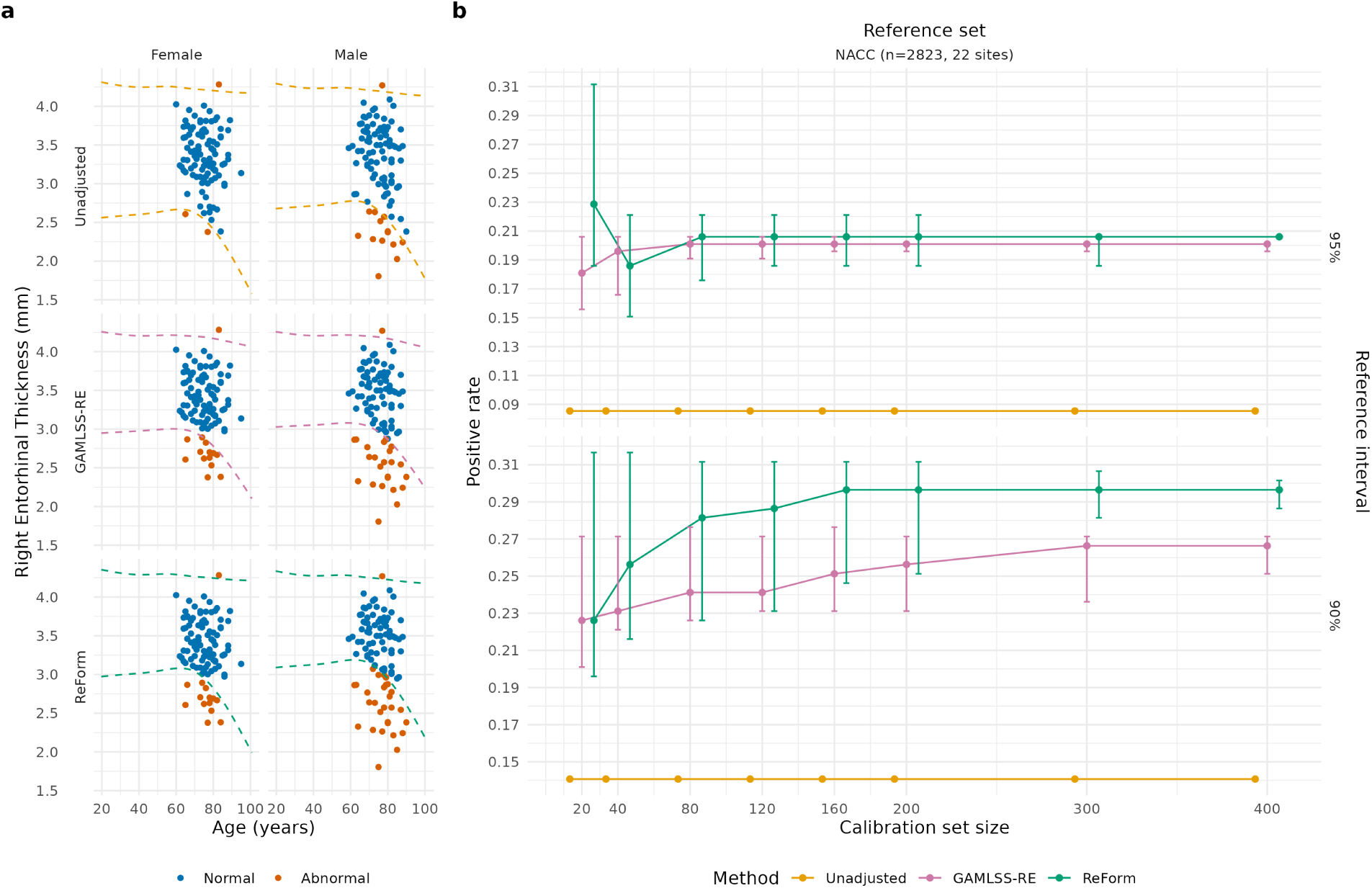
Application of reference intervals fitted in the NACC reference dataset to OASIS 3.0 T Alzheimer’s disease (AD) subjects in Experiment 3, where we fitted reference intervals in multi-site NACC healthy individuals and applied them to OASIS 3.0 T Alzheimer’s disease patients. (a) Illustration of methods using a single trial from Experiment 3. 199 OASIS 3.0 T AD scans are shown. Direct application of 95% reference intervals (Unadjusted) yielded 17 outside (8.5%). A 95% reference interval constructed using GAMLSS with a random effect across the 2928 NACC scans and 200 OASIS 3.0 T healthy scans yielded 38 outside (19.1%). A ReFormed reference interval yielded 44 outside (22.1%) (b) Positive rates across unadjusted, ComBatLS, GAMLSS-RE and ReForm for Experiment 3. Median and interquartile range are plotted across 1,000 trials. Unadjusted represents direct application of reference intervals fit using GAMLSS. Here, calibration sets are drawn from the OASIS 3.0 T cognitively normal individuals. ComBatLS is a harmonization method that uses the reference data and calibration set to adjust the data. GAMLSS-RE refers to fitting a GAMLSS with random effect for study site across the combined reference and calibration sets. ReForm is our proposed method which calibrates the unadjusted reference interval. All positive rate values are computed in the same 199 OASIS 3.0 T AD scans.

## 4 Discussion

We have introduced ReForm, a new method for calibration of reference intervals, and demonstrated its utility for MRI studies. As an illustrative example, we have focused on right entorhinal cortical thickness which is known to be lower in AD patients, and showed that ReForm considerably outperforms unadjusted intervals, refitting, and harmonization. When multi-site reference data was available, ReForm performed similarly to GAMLSS-based model adjustment for sufficiently sized samples. In application to AD patients, we show that ReForm and GAMLSS model adjustment increased the positive rate, defined as the proportion of AD patients identified as abnormal. ReForm achieved slightly higher positive rates depending on the chosen reference interval. While we compared ReForm to methods that utilize the reference dataset, ReForm does not require any reference data. Not requiring reference data is of high importance in practice due to privacy concerns related to sharing patient data as well as computational concerns if models must be refit on reference data. Moreoever, across our experiments, ReForm performance did not depend on the reference set size, model choice, or underlying data distribution—–unlike other strategies. Yet, ReForm outperformed harmonization methods and performs comparably to a GAMLSS model adjustment method, both of which require using the entire reference dataset for estimation.

Across experiments, ReForm performance depended on the calibration set size. In general, large calibration sets (≥ 120) were necessary to achieve no significant difference from the target level *α* for a given reference interval. For adequate performance, we found that about 40 samples were necessary for 90% reference intervals and 80 samples for 95% reference intervals. For 95% and 99% reference intervals across all experiments, we observed that ReForm was significantly overconservative in calibration sets of size 300. This phenomenon may be related to rounding in the computation of empirical quantiles used in ReForm. Future studies should consider interpolation as a potential solution to this issue. In Experiment 3, we observed in 99% reference intervals calibrated with less than 200 samples that ReForm exhibited poor control of FPR. For a calibration set size of less than 200, ReForm adjusts the upper bound to the largest observation and lower bound to the smallest observation. In this setting, ReForm can thus exhibit dramatically different behavior depending on outlier observations in the calibration set. For 99% reference intervals, we generally found that ReForm showed good control of FPR in calibration set sizes of 120 or greater, but advise caution due to the large variability. These findings are consistent with previous studies of GAMLSS and additive quantile regression, which suggest that large samples are necessary to estimate more extreme quantiles (Bozek et al., 2023; Jennen-Steinmetz, 2014). Overall, we suggest that the calibration set size and observations are both carefully chosen to minimize issues with FPR using ReForm. Future work could also consider other methods for conformal prediction under distribution shift, which offer solutions when less or no observations are available from the new sample (Barber et al., 2023; Ai and Ren, 2025).

While refitting is recommended by the CLSI when reference intervals fail to be verified in a new sample (Clinical and Laboratory Standards Institute, 2008), our results suggest that refitting is rarely the best option. Refitting was particularly prone to error in smaller sample sizes, and led to inflated FPR in several settings. Furthermore, refitting was dependent on the choice of model, which we demonstrated by comparing additive quantile regression versus GAMLSS. Refitting can also lead to reference intervals with unreasonable age trends as illustrated in Figure 1c. Nonetheless, in cases where no reference data or reference intervals are available, refitting shows good control of FPR with sample sizes of 160 or above. These findings are consistent with the CLSI guidelines for fitting nonparametric 95% reference intervals, which recommend at least 120 samples (Clinical and Laboratory Standards Institute, 2008).

When a multi-site reference dataset is available, model adjustment provides a solution with reasonable control of FPR. In fact, model adjustment using the entire reference dataset outperformed ReForm in smaller samples. Our result is consistent with recent studies which release GAMLSS models accompanied by out-of-sample fitting of random effects, showing generally good performance with at least 100 samples (Bethlehem et al., 2022). Our current work only considers GAMLSS model adjustment using the full reference dataset, which represents the best possible scenario for model adjustment. Future studies should evaluate how limited access to the reference dataset impacts performance of model adjustment for controlling FPR in reference intervals.

Compared to harmonization, ReForm outperformed both ComBat-GAM and ComBatLS for control of FPR across experiments with adequate calibration set size. Both ComBat-GAM and ComBatLS adjust for batch effects in mean and variance, while accounting for covariate effects. As a result, neither method explicitly models the tails of the distribution, which drive the constructed reference intervals. Comparing ComBat-GAM versus ComBatLS, we observed that ComBatLS performs substantially better, likely due to its modeling of covariate effects in the variance. We also observed that these harmonization methods can be either overly conservative or liberal in terms of FPR depending on the experimental setting. These harmonization methods were also highly dependent on the size of the reference dataset provided, which is important to consider for studies that share reference data (Zhu et al., 2025).

Comparing Experiments 1 and 2, where Experiment 2 has a lesser difference between the reference and test datasets, we observed a counterintuitive result in our harmonization methods considered. As hypothesized, Experiment 1 demonstrated poorer FPR control in unadjusted intervals compared to Experiment 2. ComBatLS offered notable benefit over unadjusted intervals in Experiment 1, but almost no benefit in Experiment 2. Several potential reasons should be considered. First, as pointed out previously, ComBatLS adjusts for batch effects in mean and variance and does not necessarily adjust the tails of the distribution used in reference interval construction. Second, minor batch effects in the mean and variance lead to minor harmonization adjustments overall. Thus, it is possible that the batch effects observed in Experiment 2 do not substantially impact the mean and variance of measurements, leading to insufficient adjustment by ComBatLS.

Several limitations of the current work limit the generalizability of our findings. First, we only considered a single MRI phenotype for its relevance to AD research. Second, our experiments involved a small number of MRI studies and cannot directly generalize to other MRI studies. While ReForm makes no assumptions about the data distribution, the demographic and phenotypical variances may impact empirical performance. Further studies should assess how ReForm across other MRI phenotypes and studies, including volumetric phenotypes and MRI studies collecting functional and diffusion MRI.

We compared ReForm to alternative methods on the basis of FPR, not on the basis of recovering the true underlying population quantiles. Our reasoning was that ReForm is a way to calibrate existing reference intervals, not construct new ones from the reference dataset. As such, we did not aim to perform a comprehensive comparison across modeling methods. Prior work has provided comparisons across methods for building brain charts in both collected MRI data (Ge et al., 2024) and simulated data (Bozek et al., 2023), as well as demonstrations of conformal prediction for controlling FPR in simulation (Lei et al., 2018; Romano et al., 2019b). Future studies could examine how well ReForm recovers true population quantiles by simulating different reference and test dataset distributions.

Our alternative methods were chosen on the basis of potential feasibility in the same setting as ReForm. Through recent papers, approximations of the statistical harmonization and GAMLSS model adjustment methods in our study can be implemented without access to the reference dataset (Bethlehem et al., 2022; Xin et al., 2025). However, our study applied those methods with access to the reference to represent the best possible performance of the alternative methods. Other harmonization methods including deep learning approaches were not implemented since our data only includes derived MRI phenotypes rather than the full MRI image (Hu et al., 2023). We chose not to compare to the ConQuR harmonization method since it does not have an implementation for out-of-sample scans (Ling et al., 2022). Our goal was to compare to a subset of prominent model frameworks while acknowledging that there are likely other approaches that could be utilized in this setting. Future studies could compare ReForm to additional methods.

Throughout this study, we evaluated FPR across all ages and sexes in the validation sample. This FPR measure has limitations though, since it does not guarantee that reference intervals control FPR within age and sex bins. An extension of conformal prediction suggests that sampling calibration sets across groups of interest is a viable solution to this problem (Romano et al., 2019a). Other recent works also consider this problem and propose extensions of conformal prediction that guarantee coverage conditional on covariates (Gibbs et al., 2023; Guan, 2023; Hore and Barber, 2024). Further investigation is needed to determine if their methodologies can be applied in our setting and to assess the sample size requirements and empirical performance. Follow-up studies in MRI reference intervals should consider defining appropriate age and sex bins for evaluation of FPR within demographic groups.

Finally, we advise caution in interpreting MRI reference intervals. To quote Leo Tolstoy’s *Anna Karenina*, ”[a]ll happy families are alike; each unhappy family is unhappy in its own way”. In other words, an observation that falls outside an MRI reference interval may be abnormal owing to a multitude of reasons. In this work, we do not investigate potential causes. In practice, it is important to treat an abnormal observation with particular attention and closely examine other measurements to understand potential biological or technical causes.

## 5 Conclusion

We propose a novel method called ReForm which calibrates reference intervals to a new sample. Unlike other methods, we show that the performance of ReForm does not depend on the size of the reference dataset or nature of the modeling approach used to fit reference intervals. If a large multi-site reference dataset is available, then model-based batch effect adjustment offers a similarly viable solution that performs well even in small sample sizes. Our recommendations are to consider ReForm across a wide range of settings, so long as the sample size is sufficient for type of reference interval considered. We find that 40 samples is adequate for 90% reference intervals and 80 samples for 95% reference intervals. For the challenging problem of 99% reference intervals, we find that ReForm requires at least 120 samples to ensure acceptable control of FPR, but advise caution due to sensitivity to outliers. These recommendations are based on a limited set of MRI studies and benchmarks, which should be expanded in future work. While we have focused on a single MRI phenotype in our study, our ReForm method is broadly applicable to any phenotype that differs across study sites.

## Data and Code Availability

An R implementation of the proposed method, ReForm, is available via GitHub (https://github.com/andy1764/ReForm). Data used in this work were from the Lifespan Brain Chart Consortium (LBCC), which includes multiple publicly-available MRI datasets. Our work included three datasets, each with their own data access and sharing policies, which can be accessed via the following websites: The Alzheimer’s Disease Repository Without Borders (ARWiBo, https://www.arwibo.it); Open Access Series of Imaging Studies (OASIS, https://sites.wustl.edu/oasisbrains/); and the National Alzheimer’s Coordinating Center (NACC, https://naccdata.org/).

## Author Contributions

A.A.C.: Conceptualization, Methodology, Software, Validation, Formal Analysis, Investigation, Resources, Data Curation, Writing—Original Draft, Writing—Review and Editing, Visualization, Funding Acquisition. J.S.: Resources, Data Curation, Writing—Review and Editing. Resources, Data Curation, Writing—Review and Editing. M.G.: Resources, Data Curation, Writing—Review and Editing. R.A.I.B: Resources, Data Curation, Writing—Review and Editing. L.D.: Resources, Data Curation, Writing—Review and Editing. E.K.: Resources, Data Curation, Writing—Review and Editing. A.B.: Writing—Review and Editing. J.H.J.: Writ-ing—Review and Editing. S.V.: Methodology, Writing—Review and Editing. T.D.S.: Writing—Review and Editing. A.A.B.: Conceptualization, Resources, Data Curation, Writing—Original Draft, Writing—Review and Editing, Visualization, Supervision, Funding Acquisition.

## Ethics

All subjects gave written informed consent prior to participation and all studies were approved by the regional ethics committees.

## Acknowledgments

This work was supported by the National Institute of Mental Health (award numbers R01MH133843, R01MH123550, R01MH112847, R01MH113550, R37MH125829, and R01MH120482). A.B. and J.H.J. were supported by the National Institutes of Health (R01AG054159); this grant provided effort support and did not fund the data collection or analyses reported in this manuscript. The content is solely the responsibility of the authors and does not necessarily represent the official views of the National Institutes of Health.

