## Supplementary Materials for "Calibration of MRI-based reference intervals to new samples"

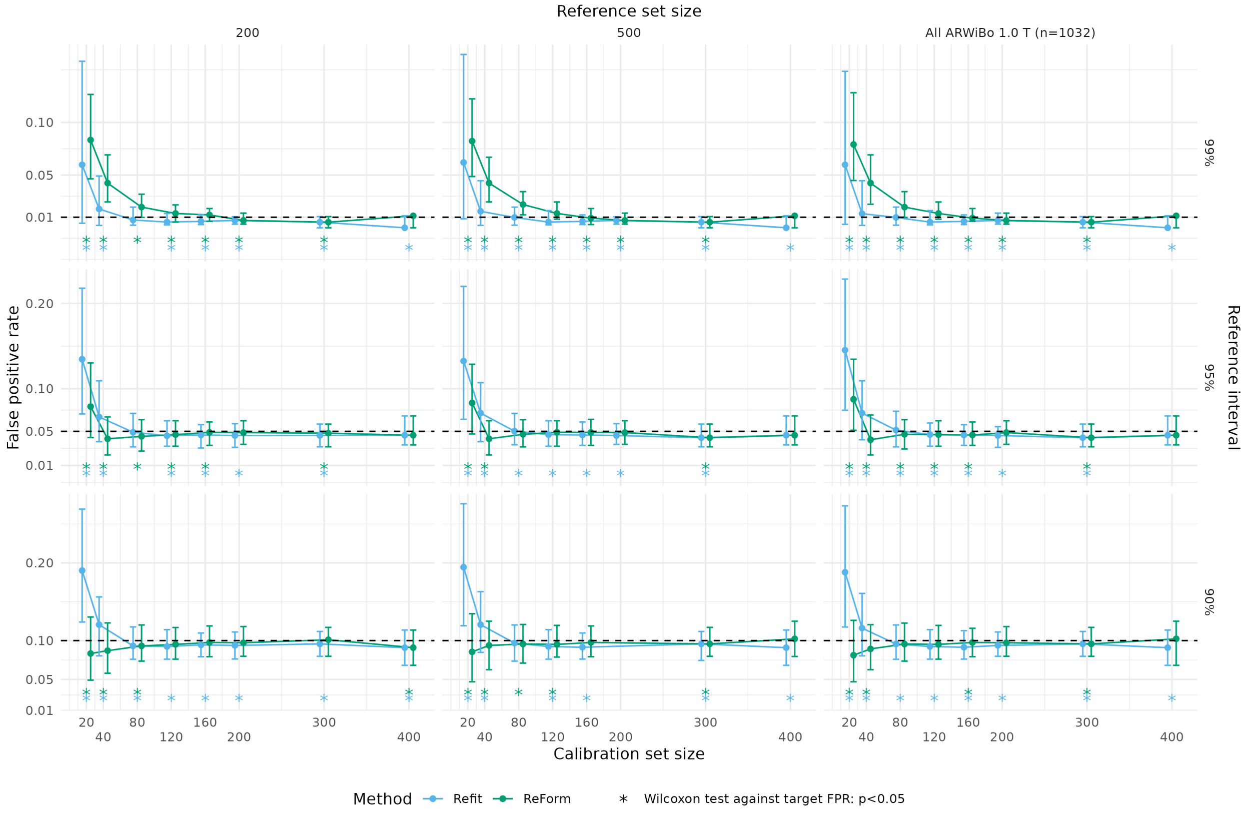


Figure 1: False positive rates across unadjusted, refit, and ReFormed reference intervals from Experiment 1, which fitted reference intervals using additive quantile regression in ARWiBO 1.0 T scans and applied them to OASIS 3.0 T scans. Median and interquartile range are plotted across 1,000 trials. Plots assess the validity of estimated 99% reference intervals (top panel, target FPR of 0.01); 95% reference intervals (middle panel, target FPR of 0.05; and 90% reference interval (bottom panel, target FPR of 0.1). Asterisks used to show significant differences from the target FPR (dashed line) using a one-sample Wilcoxon signed-rank test. Unadjusted represents direct application of reference intervals fit using additive quantile regression. Refit refers to refitting using additive quantile regression in the calibration set of a designated size. ReForm is our proposed method which calibrates the unadjusted reference interval. All FPR values are computed in the same OASIS 3.0 T scans with varying sample size left after holding out the calibration set. In Refit, some trials yielded errors during fitting and are not shown.

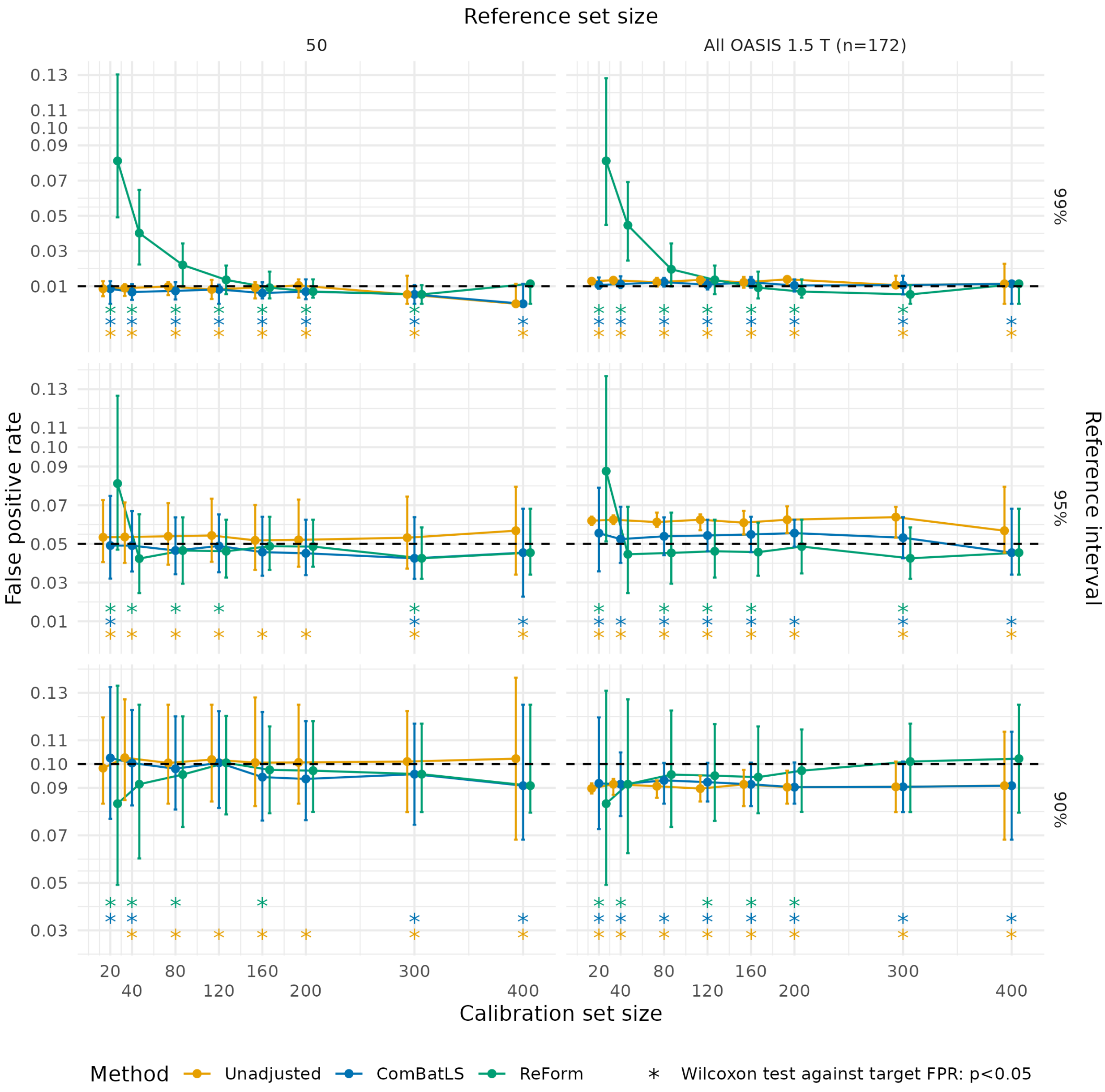


Figure 2: False positive rates across harmonization methods and ReForm from Experiment 1, which fitted reference intervals using additive quantile regression in ARWiBO 1.0 T scans and applied them to OASIS 3.0 T scans. Median and interquartile range are plotted across 1,000 trials. Plots assess the validity of estimated 99% reference intervals (top panel, target FPR of 0.01); 95% reference intervals (middle panel, target FPR of 0.05; and 90% reference interval (bottom panel, target FPR of 0.1). Asterisks used to show significant differences from the target FPR (dashed line) using a one-sample Wilcoxon signed-rank test. ComBat-GAM and ComBatLS are fit on the entire reference set combined with the calibration set. ReForm is our proposed method which calibrates the unadjusted reference interval. All FPR values are computed in the same OASIS 3.0 T scans with varying sample size left after holding out the calibration set.

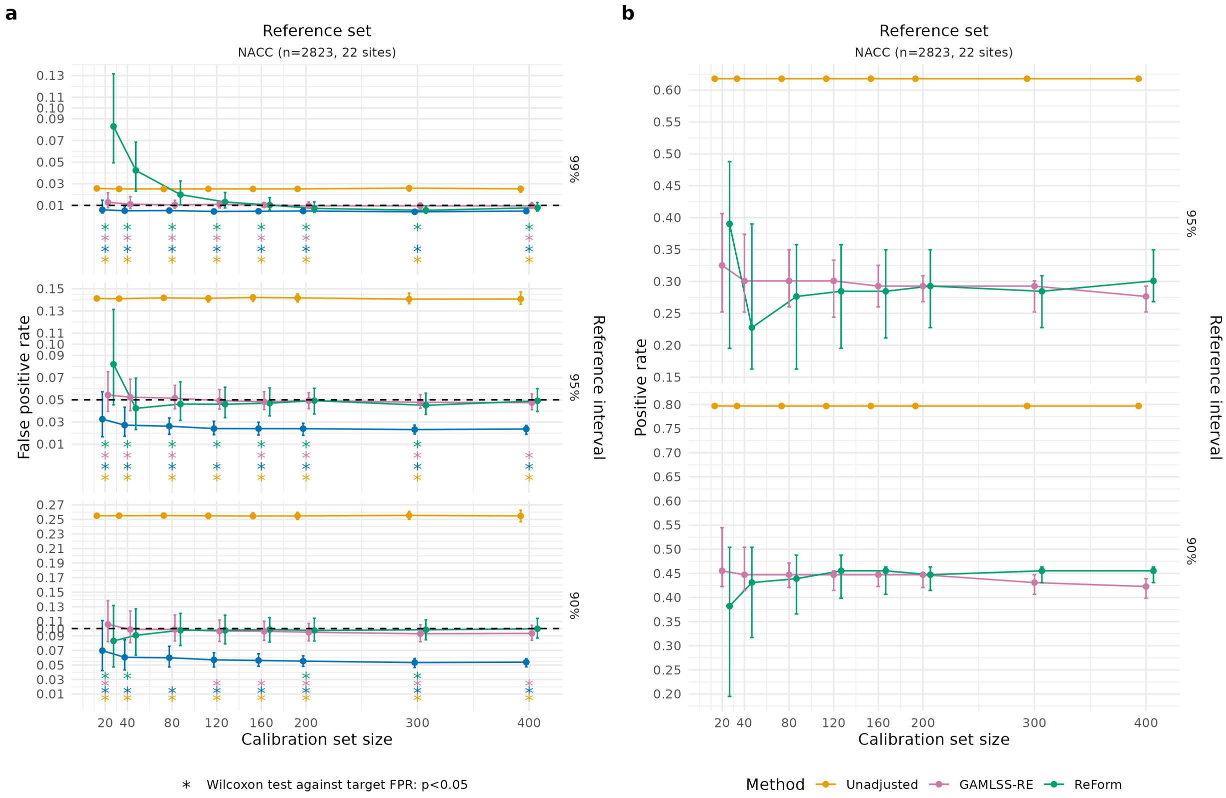


Figure 3: Application of reference intervals fitted in the NACC reference dataset to ARWiBo 1.0 T healthy (a) and Alzheimer's disease (AD) subjects (b). (a) False positive rates across unadjusted, ComBatLS, GAMLSS-RE and ReForm for fitting reference intervals in NACC scans and applying to ARWiBo healthy scans. Median and interquartile range are plotted across 1,000 trials. Plots assess the validity of estimated 99% reference intervals (top panel, target FPR of 0.01); 95% reference intervals (middle panel, target FPR of 0.05; and 90% reference interval (bottom panel, target FPR of 0.1). Asterisks used to show significant differences from the target FPR (dashed line) using a one-sample Wilcoxon signed-rank test. (b) Positive rates within this subset of trials across unadjusted, GAMLSS-RE and ReForm for fitting reference intervals in NACC healthy individuals and applying to ARWiBo AD subjects. Median and interquartile range are plotted across 1,000 trials. Unadjusted represents direct application of reference intervals fit using GAMLSS. ComBatLS is fit on the entire reference set combined with the calibration set. GAMLSS-RE refers to fitting a GAMLSS with random effect for study site across the combined reference and calibration sets. ReForm is our proposed method which calibrates the unadjusted reference interval. All positive rate values are computed in the same 123 ARWiBo AD scans.

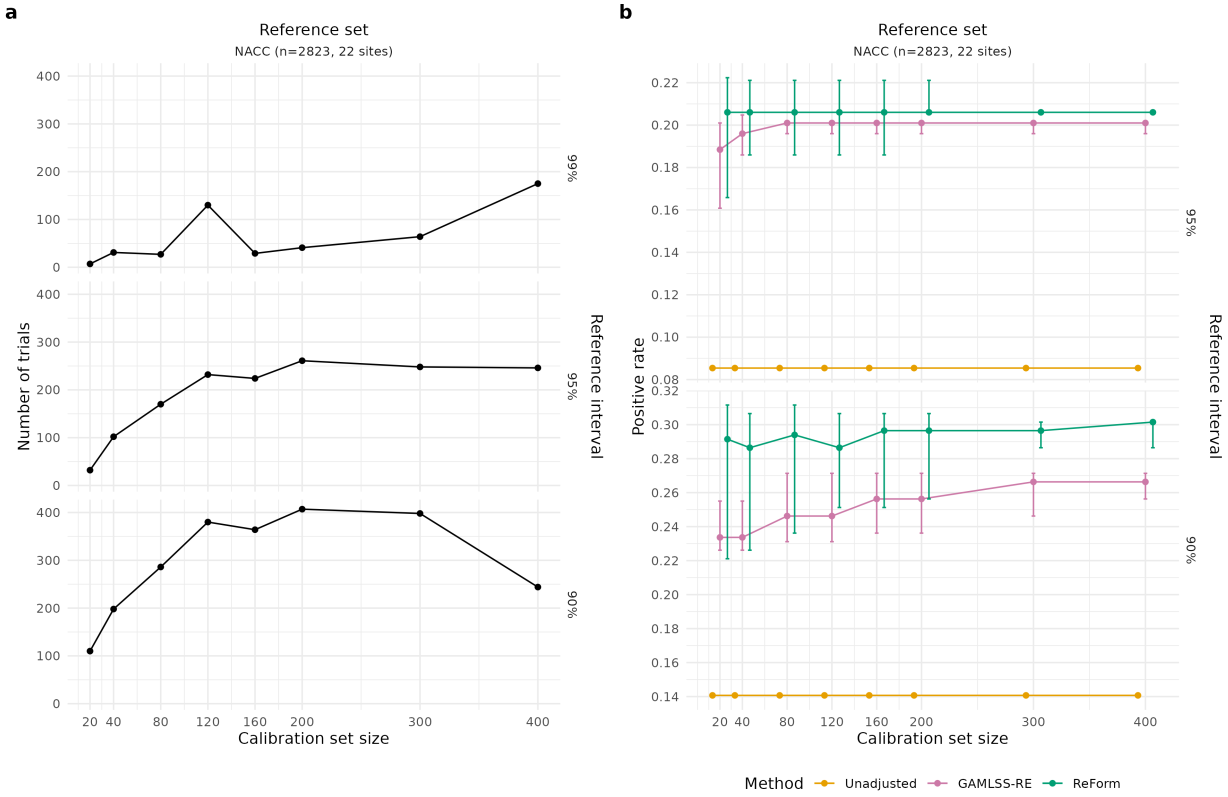


Figure 4: Application of reference intervals fitted in the NACC reference dataset to OASIS 3.0 T Alzheimer's disease (AD) subjects including only trials with controlled false positive rate. (a) Number of trials where both GAMLSS with random effects (GAMLSS-RE) and ReForm achieve false positive rate within OASIS 3.0 T cognitively normal scans near the target FPR ($\alpha$) with a tolerance of $0.2 \times \alpha$. (b) Positive rates within this subset of trials across GAMLSS-RE and ReForm for fitting reference intervals in NACC healthy individuals and applying to OASIS 3.0 T Alzheimer's disease (AD) subjects. Median and interquartile range are plotted across 1,000 trials. Unadjusted represents direct application of reference intervals fit using GAMLSS. GAMLSS-RE refers to fitting a GAMLSS with random effect for study site across the combined reference and calibration sets. ReForm is our proposed method which calibrates the unadjusted reference interval. All positive rate values are computed in the same 319 OASIS 3.0 T AD scans.

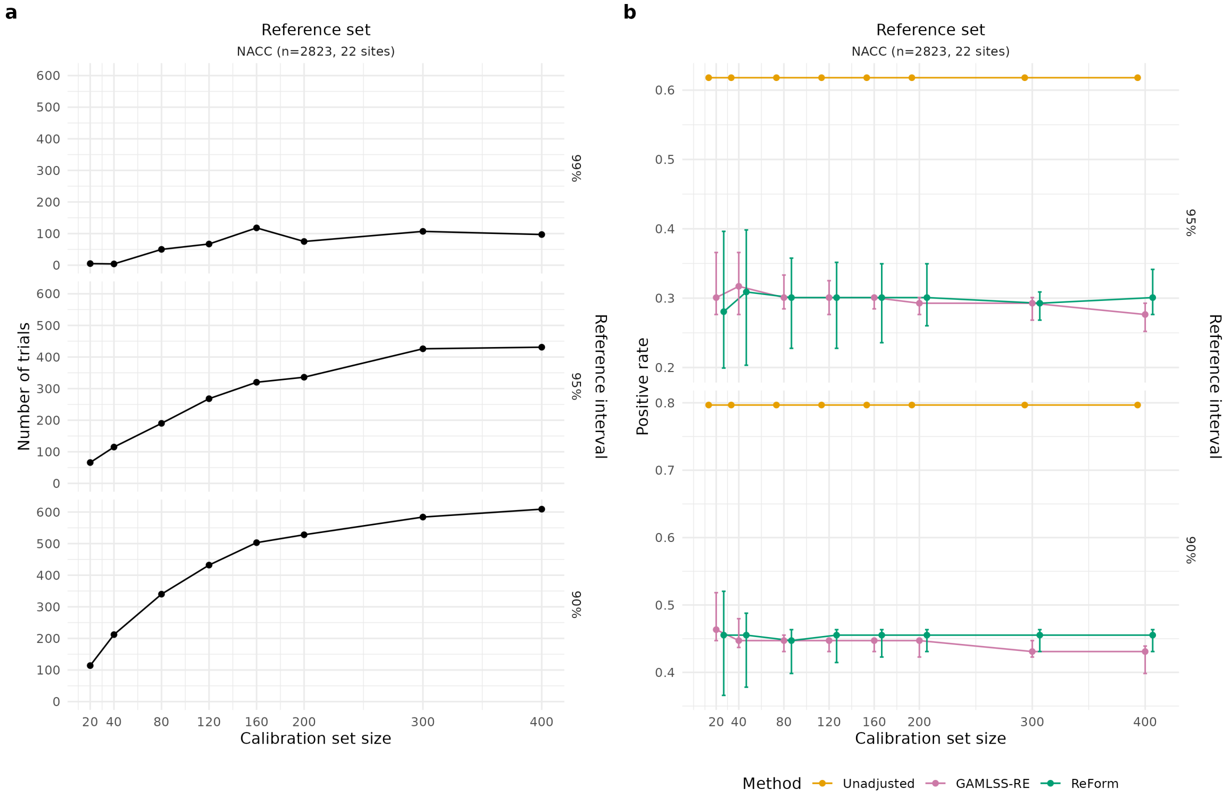


Figure 5: Application of reference intervals fitted in the NACC reference dataset to ARWiBo 1.0 T Alzheimer's disease (AD) subjects including only trials with controlled false positive rate. (a) Number of trials where both GAMLSS with random effects (GAMLSS-RE) and ReForm achieve false positive rate within ARWiBo cognitively normal scans near the target FPR ($\alpha$) with a tolerance of $0.2 \times \alpha$. (b) Positive rates within this subset of trials across GAMLSS-RE and ReForm for fitting reference intervals in NACC healthy individuals and applying to ARWiBo Alzheimer's disease (AD) subjects. Median and interquartile range are plotted across 1,000 trials. Here, calibration sets are drawn from the ARWiBo cognitively normal individuals. GAMLSS-RE refers to fitting a GAMLSS with random effect for study site across the combined reference and calibration sets. ReForm is our proposed method which calibrates the unadjusted reference interval. All positive rate values are computed in the same 123 ARWiBo AD scans.
